# Development and characterization of influenza M2 ectodomain and/or HA stalk-based DC-targeting vaccines for different influenza infections

**DOI:** 10.1101/2021.10.07.463539

**Authors:** Titus Abiola Olukitibi, Zhujun Ao, Hiva Azizi, Mona Mahmoudi, Kevin Coombs, Darwyn Kobasa, Gary Kobinger, Xiaojian Yao

## Abstract

A universal influenza vaccine is required for broad protection against influenza infection. Here, we revealed the efficacy of novel influenza vaccine candidates based on Ebola glycoprotein (EboGP) DC-targeting domain (EΔM) fusion protein technology. We fused influenza hemagglutinin stalk (HAcs) and extracellular matrix protein (M2e) or four copies of M2e (referred to as tetra M2e (tM2e)) with EΔM to generate EΔM-HM2e or EΔM-tM2e, respectively, and revealed that EΔM facilitates DC/macrophage targeting *in vitro*. In a mouse study, EΔM-HM2e- or EΔM-tM2e-pseudotyped viral particles (PVPs) induced significantly higher titers of anti-HA and/or anti-M2e antibodies. We also developed recombinant vesicular stomatitis virus (rVSV)-EΔM-HM2e and rVSV-EΔM-tM2e vaccines that resulted in rapid and potent induction of HA and/or M2 antibodies in mouse sera and mucosa. Importantly, vaccination protects mice from influenza H1N1 and H3N2 challenges. Taken together, our study suggests that recombinant rVSV-EΔM-HM2e and rVSV-EΔM-tM2e are efficacious and protective universal vaccines against influenza.

---

Influenza is a highly contagious airborne disease that attacks the respiratory system and occurs in seasonal epidemics and pandemics. The influenza pandemic in 1918 killed approximately 50 million people globally ^1, 2^, and to date, influenza virus infection is still posing a substantial threat to the health sector worldwide ^3^. Circulating influenza vaccines are associated with some issues, including the level of effectiveness of annual vaccines protecting against the specific influenza strain in the particular seasonal epidemic and the psychological effect on the population who must receive a flu shot every year for their lifetime. Based on these findings, the Centers for Disease Control (CDC), as part of their recent recommendations, emphasizes a need for a universal vaccine against influenza viral infection^4^.

The universal vaccine is characterized by the ability to protect individuals from different strains of the influenza virus. In addition to the four different types of influenza, each type is composed of a population of different strains. Variation in influenza strains occurs at hemagglutinin (HA) and neuraminidase (NA). For instance, influenza A, which is the most prominent family, has eighteen known HA subtypes and eleven identified NA subtypes with different host ranges, including humans, birds, bats, and swine ^4, 5^. The difficulty in producing a universal vaccine against the influenza virus is due to the antigenic shift or antigenic drift caused by reassortment or mutation of HA or NA. These mechanisms allow the influenza virus to continuously escape host immune defenses ^6^. Additionally, the current vaccine in circulation is short-lived and narrow^7^. Therefore, as a method to develop a universal vaccine, the conserved components on the surface protein of influenza could be utilized to elicit broad immune responses specific for all the present and future strains of the influenza virus.

The influenza HA protein, which is responsible for the cell attachment and entry of the viruses, is a promising epitope in the development of influenza vaccines. Although the globular head of HA induces the production of neutralizing antibodies ^8, 9^, it is difficult to use when developing a universal vaccine due to the large number of HA subtype variations. However, the HA stalk and the highly conserved extracellular matrix protein (M2e) have been found to be promising in the development of a universal vaccine for influenza viral infection due to their durability and stability ^10, 11^ These subunit proteins of influenza induce broader neutralization and participate in either antibody-dependent cellular phagocytosis (ADCP) or antibody-dependent cellular cytotoxicity (ADCC), which subsequently eliminate influenza virus or destroy the cells already infected ^12, 13^. Numerous studies have used different approaches to develop universal influenza vaccines, including fusion of influenza M2e polypeptides ^14^, targeting conserved broadly reactive epitopes on the HA stalk ^15^, developing fusion proteins between influenza M2e and bacterial flagellin ^16^, expressing recombinant HA in virus-like particles ^17^ and using VSV to deliver HA antigens ^18^. However, although some of these approaches are being investigated in clinical trials ^19^, varying limitations still exist, with the majority having relatively low immune responses, except for VSV that is used with adjuvants such as MF59 and ASO3 ^6^. Therefore, new and efficient universal vaccine(s) with broad protection against various strains of the influenza virus must be developed.

The dendritic cell (DC)-targeting vaccine has recently received global attention since this approach is effective because DCs function as antigen-presenting cells (APCs) that stimulate adaptive immune responses and regulate innate immune responses ^20 21^. The usage of this technology is in the pipeline for the development of various vaccines against viral pathogens and cancers^20 21 22^. A study showed that targeting influenza HA and chemokine receptor Xcr1^+^ to DCs induces immune responses and confers protection against the influenza virus ^23^. A group of scientists also targeted influenza M2e to DCs by fusing M2e with anti-Clec9 ^24^, while HA of influenza was infused with an artificial adjuvant vector cell targeting DCs to induce CD4^+^ T cells and CD4^+^ Tfh cells ^25^.

Ebola virus glycoprotein (EboGP) is the viral protein expressed on the Ebola virus (EBOV) surface that preferentially binds to DCs, monocytes and macrophages ^26, 27^. As shown in our recent study, the incorporation of EboGP into HIV PVPs indeed facilitates DC and macrophage targeting and significantly enhances HIV-specific immune responses ^28, 29^. These observations indicated that EboGP has the potential to direct an HIV antigen toward DCs to facilitate effective anti-HIV immune responses. Notably, a highly glycosylated mucin-like domain (MLD) encompassing residues 313 to 501 is located at the apex and the sides of each EboGP monomer ^30^. However, some studies have shown that removing this MLD region does not impede EboGP-mediated lentiviral vector entry ^31^ and was dispensable for EBOV infections *in vitro* ^32, 33^. Our laboratory has recently developed the EboGP mucin-like domain replacement system and shown the great potential of this vaccine technology to deliver heterologous polypeptides *in vivo* and stimulate innate and adaptive immune responses ^34^.

In this study, we used this EboGP DC-targeting domain-based fusion protein technology to fuse conserved HA stalk regions (HAcs) and M2e or four copies of M2e (referred to as tetra M2e (tM2e)) within the MLD-deleted EboGP (EboGPΔM). By incorporating these fusion proteins into HIV-based pseudotyped viral particles (PVPs) or a recombinant vesicular stomatitis virus (rVSV) vector, we characterized their DC-targeting ability and investigated their potential for eliciting host immune responses and their abilities to protect against H1N1 and H3N2 influenza infections.

## Results

### 1. Construction and characterization of EΔM-tM2e and EΔM-HM2e chimeric fusion proteins

The HAcs and M2e proteins of the influenza virus are highly conserved among the strains of the influenza virus, and previous studies have attempted to develop HAcs- or M2e-based universal vaccines for influenza virus ^8, 9, 10, 11, 35^. We used the EboGPΔM Replacement System ^34^ to insert 4 copies of M2e (tM2e) (aa 92) into EΔM and form the EΔM-tM2e fusion protein as a method to improve the DC-targeting ability of HAcs- or M2e-based vaccines (Fig. 1A, C). The four copies of influenza M2e consisted of M2e from human (2 copies), swine (1 copy) and bird (1 copy) strains and were linked with a GGGS linker (Fig. 1A). Additionally, we combined a copy of M2e from the human influenza strain with the HAcs from influenza H5N1 using a GSA linker (HM2e) (aa 179) and inserted it into EΔM to form EΔM-HM2e as a fusion protein (Fig. 1B, C).

**Figure 1:**
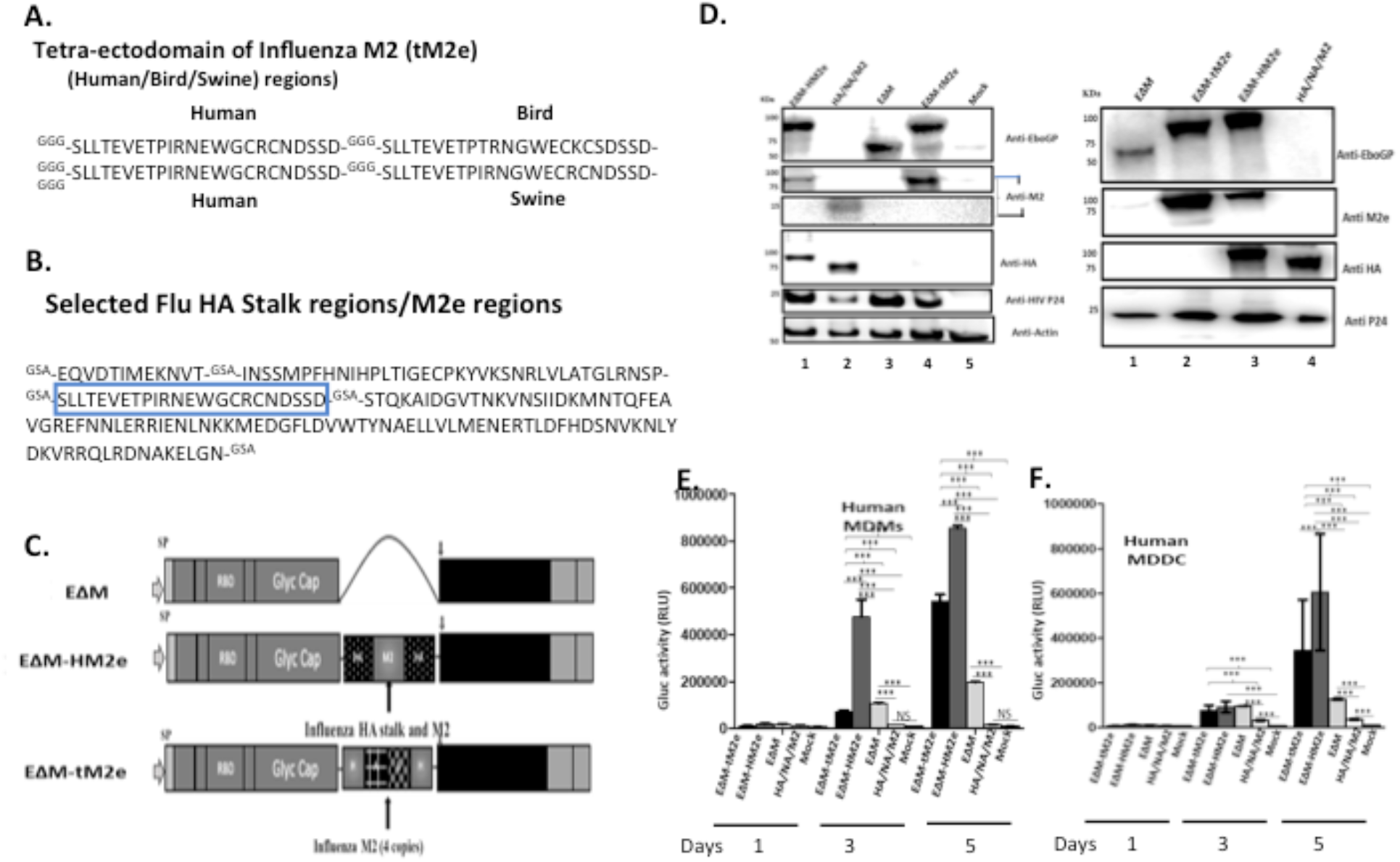
The construction, expression and cell entry ability of EΔM-tM2e or EΔM-HM2e. A) Amino acid sequences of tM2e including influenza virus M2e consensus from human, swine, and bird strains. B) Amino acid sequence of HM2e including influenza virus HA stalk and M2e from human strain. C) Codon-optimized EboGP gene with MLD deletion (aa 305 – 485)(EΔM) (upper lane) and pCAGGS-EΔM-HM2e or pCAGGS-EΔM-tM2e, respectively. D) 293T cells were co-transfected pCAGGS-EΔM-tM2e, pCAGGS-EΔM-HM2e or pCAGGS-HA/NA/M2, together with Δ8.2 and ΔRI/ΔE/Gluc+. WB was used to detect the expression of M2e, HA, EboGP and HIV P24 in cells (left panel) and on VPVs (right panel). E-F) Human PBMC derived macrophages (MDMs), and MDDCs were infected with the EΔM-HM2e, EΔM-tM2e, EΔM and native HA/NA/M2-PVPs. The supernatants were collected at different times of infection and subjected to a GLuc activity assay. Error bars represent variation between duplicate samples, and the data are representative of results obtained in three independent experiments.

We first produced PVPs that incorporated the EΔM-tM2e or EΔM-HM2e fusion protein and investigated their abilities into enter DCs/macrophages. Briefly, EΔM-tM2e- or EΔM-HM2e-expressing plasmids were cotransfected with an HIV Gag-Pol packaging plasmid Δ8.2 and a multigene (reverse transcriptase/integrase/envelope)-deleted HIV vector (ΔRI/ΔE/Gluc) in HEK-293T cells as previously described ^28, 36^. In ΔRI/ΔE/Gluc, the *nef* gene was replaced with a Gaussia luciferase (Gluc) gene that was expressed and released when the virus entered cells. Meanwhile, EΔM- and native influenza HA/NA/M2-pseudotyped PVPs ^37^ were included as controls. Forty-eight hours after transfection, the cells and supernatants containing PVPs were collected, and the presence of different proteins in cells and PVPs was assessed using Western blotting (WB) with the corresponding antibodies (Fig. 1D and E). As expected, the anti-EboGP antibody detected the expression of EΔM, EΔM-HM2e or EΔM-tM2e fusion proteins (Fig. 1D, upper panel, Lanes 1, 3, and 4). Additionally, M2e in EΔM-HM2e or EΔM-tM2e was readily detected in transfected cells using an anti-M2e antibody (Fig. 1D, second panel, Lanes 1 and 4). We also detected the expression of M2e in the influenza HA/NA/M2 sample (Fig. 1D, third panel, Lane 2). Notably, the expression levels of M2 in EΔM-tM2e-transfected cells were significantly higher than those in EΔM-HM2e-transfected cells due to the presence of 4 copies of M2e in EΔM-tM2e, while EΔM-HM2e had only one copy. Meanwhile, we detected the presence of HM2e or HA protein in EΔM-HM2e and influenza HA/NA/M2 samples (Fig. 1D, fourth panel, Lanes 1 and 2). Consistently, we also detected the presence of the fusion proteins EΔM, EΔM-HM2e or EΔM-tM2e in PVPs (Fig. 1E, Lanes 1 to 3). The presence of HIV p24 in cells and PVPs was also detected using a rabbit anti-HIV p24 antibody (Fig. 1D and E). Overall, these results indicated that EΔM-HM2e and EΔM-tM2e fusion proteins were efficiently expressed and incorporated into PVPs.

### 2. The ability of EΔM-HM2e- or EΔM-tM2e-PVPs to enter macrophages and dendritic cells (DCs)

EboGP has been shown to play a critical role during the infection of DCs and macrophages by facilitating viral attachment, fusion, and entry ^28, 38, 39^. Therefore, we investigated whether the fusion of influenza-conserved HM2e or tM2e with EΔM would impede or affect the ability of EΔM to enter DCs and macrophages. Briefly, we infected human monocyte-derived macrophages (MDMs) or monocyte-derived DCs (MDDCs) with equal amounts (adjusted with HIV p24 levels) of EΔM-tM2e- or EΔM-HM2e-pseudotyped Gluc^+^ PVPs. In parallel, the EΔM-pseudotyped Gluc^+^ PVPs were used as controls. On different days after infection, the ability of the PVPs to enter cells was monitored by detecting the Gluc activity in the supernatant of infected cells. As expected, EΔM-tM2e- and EΔM-HM-PVPs displayed more efficient entry than EΔM-PVPs (Fig. 1 F and G), suggesting that the fusion of influenza HAcs and/or M2e with EΔM did not affect the cell entry efficiency. Overall, the fusion of HM2e or tM2e to EΔM still maintains the DC- and macrophage-targeting ability of PVPs.

### 3. EΔM-HM2e- or EΔM-tM2e-PVP immunization induced significantly higher titers of specific anti-influenza HA and M2 antibodies in mice

Since EΔM-HM2e or EΔM-tM2e PVPs significantly target DCs and macrophages, we next investigated whether EΔM-HM2e or EΔM-tM2e PVPs strongly stimulated influenza HA and M2e immune responses *in vivo*. For this experiment, we subcutaneously immunized Balb/c mice with equal amounts (100 ng HIV p24) of EΔM-HM2e-, EΔM-tM2e-PVPs, or native HA/NA/M2-PVPs. The body weight of all immunized mice was monitored at 0, 28 and 56 days, and no statistically significant differences were observed (Fig. 2A and B). On Day 63 postimmunization, sera from mice were collected as described in the Materials and Methods, and the anti-M2e and anti-HA specific humoral responses were determined using ELISAs in plates coated with M2e peptides or recombinant HA from H5N1. Influenza M2e-specific humoral immune responses were detected in mice injected with EΔM-tM2e, EΔM-HM2e, and native HA/NA/M2-PVPs (Fig.2C). Interestingly, our results revealed that EΔM-tM2e PVP immunization elicited robust production of influenza M2e-specific antibodies. Additionally, antibody titers induced in the mice immunized with EΔM-HM2e-PVPs were significantly higher than those induced by native HA/NA/M2-PVPs on Day 63 (Fig. 2C). Meanwhile, immunization with EΔM-HM2e-PVPs induced significantly higher anti-HA humoral responses than immunization with HA/NA/M2-PVPs (Fig. 2D). Collectively, these results indicate that EΔM-tM2e-PVP and EΔM-HM2e-PVP immunization resulted in significantly stronger specific humoral antibodies against influenza M2e and/or HA.

**Figure 2:**
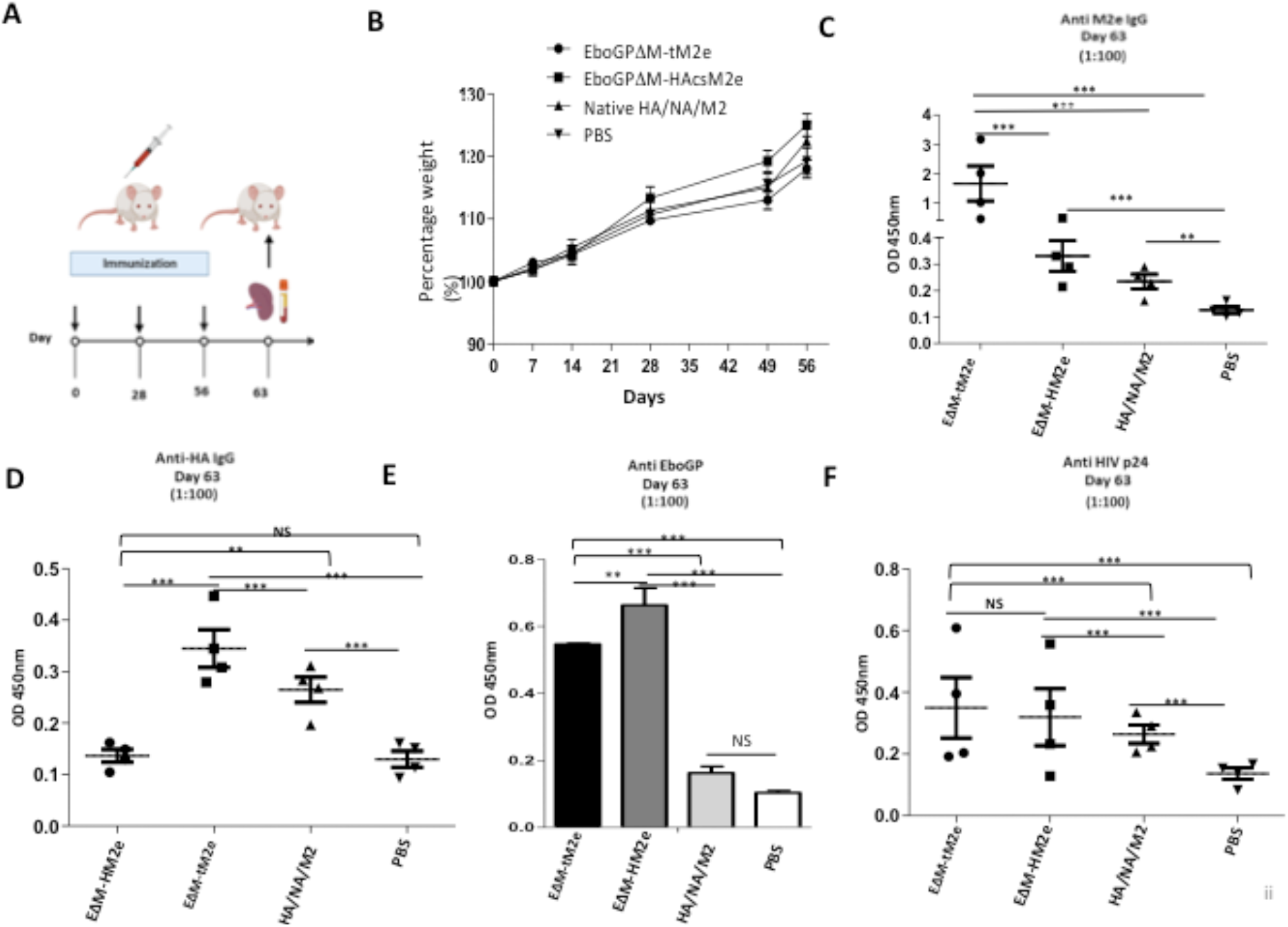
The anti- M2e and anti-HA antibodies were induced by EΔM-tM2e or EΔM-HM2e -PVPs in Balb/c mice. A) Schematic of the EΔM -tM2e, EΔM -HAcsM2e or Native HA/NA/M2 -PVPs immunization protocol used in this study. B) Mice body weights were monitored weekly, in which 100% body weight was set at day 0. The levels of Anti-M2e (C), anti-HA (D), anti-Ebola GP (E), and anti-HIV P24 (F) IgG specific antibodies were measured by ELISA method. Statistical significance was determined using an unpaired t-test, and significant p values were represented with asterisks *≤0.05, **≤0.01, ***≤0.001, ****≤0.0001. No significance (ns) was not shown.

Since both EΔM-tM2e-PVPs and EΔM-HM2e-PVPs contain the HIV-1 Gag protein and EbolaGP, we therefore tested the anti-EbolaGP- and anti-HIV P24-specific humoral immune responses in immunized mice. As expected, we only observed a significantly higher anti-EbolaGP antibody titer in the sera of mice immunized with EΔM-tM2e- and EΔM-HM2e-PVPs (Fig. 2E). Intriguingly, these results also revealed that EΔM-tM2e and EΔM-HM2e-PVPs induced significantly higher anti-HIVp24 humoral responses than native HA/NA/M2-PVPs, even when animals were immunized with equal amounts of PVPs, as quantified by HIVp24 levels (Fig. 2F). Based on these results, the presence of EΔM in the PVPs might have enhanced the antigenicity of the HIVp24 protein. However, some limitations are associated with PVP production, including the production time, cumbersome methodology and cost ineffectiveness^40^. We therefore further investigated whether the expression of EΔM-tM2e or EΔM-HM2e in the rVSV vector would induce strong immune responses against the influenza virus.

### 4. Construction and characterization of rVSV expressing the EΔM-tM2e or EΔM-HM2e chimeric protein

The rVSV vector represents a safe and potent vaccine development platform for stimulating both innate and adaptive immune responses ^41, 42, 43^. In this study, we further investigated whether the expression of EΔM-tM2e or EΔM-HM2e in the rVSV vectors also induced the production of anti-M2e and anti-HA antibodies, respectively, *in vivo*. For this purpose, we inserted the genes encoded by EΔM-tM2e and EΔM-HM2e into an rVSV vector to the position where the VSV-G gene sequence was located (Fig. 3A). Then, the attenuated replicating rVSV expressing either EΔM-tM2e or EΔM-HM2e, named rVSV/EΔM-tM2e or rVSV/EΔM-HM2e, was rescued in VeroE6 cells via reverse genetics technology ^44^, as described in the Materials and Methods. Each rVSV/EΔM-tM2e- or rVSV/EΔM-HM2e-infected vero-E6 cell line was collected and processed for WB analyses, and the results clearly showed the expression of EΔM-tM2e or EΔM-HM2e and VSV nucleoprotein (N) in the infected Vero E6 cells from the corresponding groups (Fig. 3B). We also detected the expression of EΔM-tM2e or EΔM-HM2e in Vero E6 cells using an immunofluorescence assay, as described in the Materials and Methods ^45^. Our results showed the expression of the M2e protein in rVSV-EΔM-tM2e- and rVSV-EΔM-HM2e-infected cells, while HA was only detected in rVSV-EΔM-HM2e-infected cells (Figure 3C).

**Figure 3:**
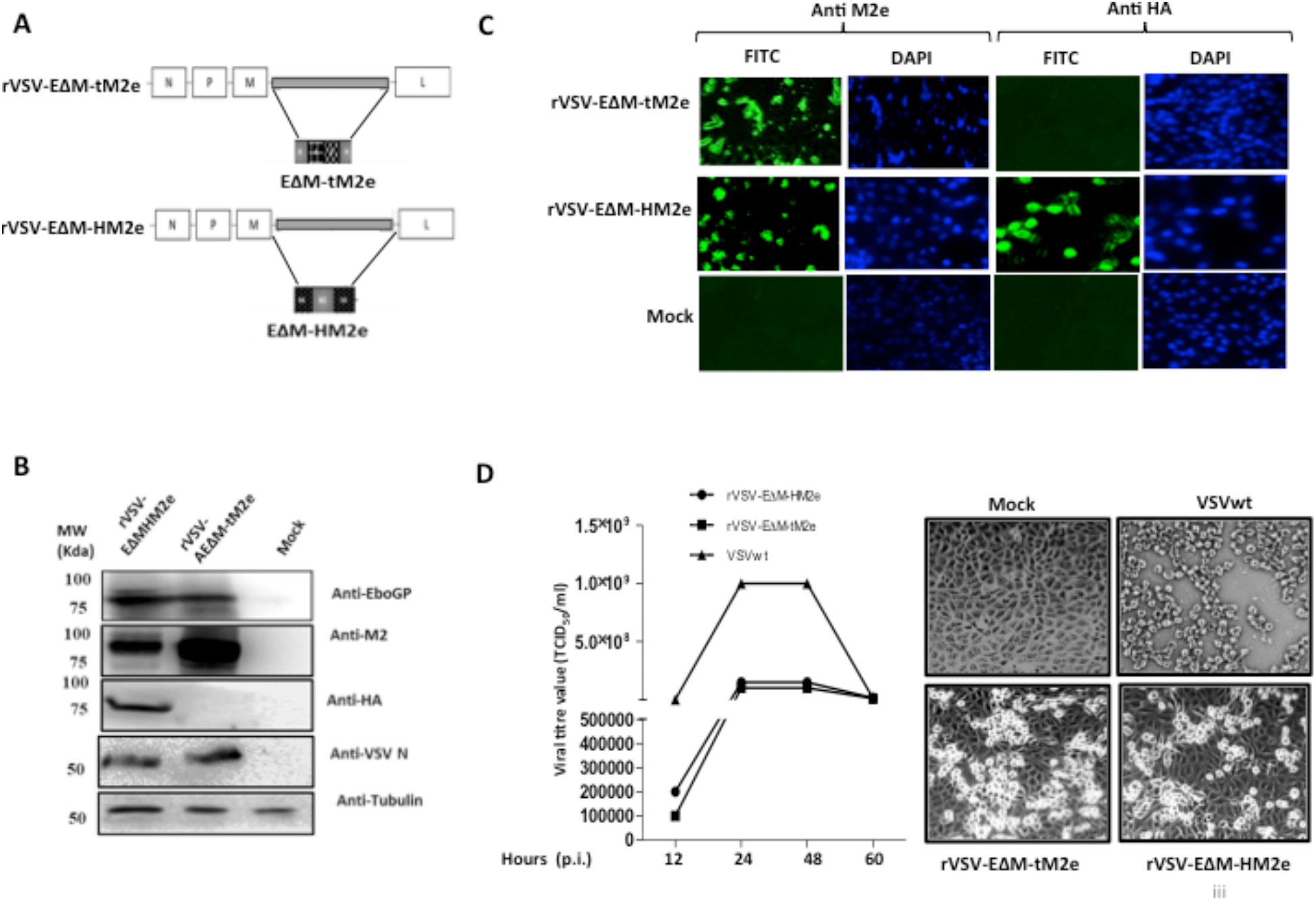
Generation and characterization of rVSV expressing the EΔM-tM2e or EΔM-HM2e. A) Schematic structures of recombinant vesicular stomatitis virus (rVSV-ΔG) vector expressing the EΔM -tM2e or EΔM-HM2e. B) The expressions of the EΔM-tM2e, EΔM-HAM2e, EboGP and VSV-N proteins in the rVSV infected Vero E6 cells by WB. C) The expression of EΔM -tM2e or EΔM-HM2e in Vero E6 cells detected by using immunofluorescence assay. D). Vero E6 cell was infected with rVSV-EΔM-tM2e, rVSV-EΔM-HM2e or VSV wt with MOI 0.001. Supernatants were collected at various time points and were titrated using Reed and Muench method^61^ (Left). At 48h post infection, the cells were observed under microscope (Right panel).

We next evaluated the replication of rVSV-EΔM-tM2e or rVSV-EΔM-HM2e in VeroE6 cells compared to wild-type rVSV (VSVwt). Vero E6 cells were infected with rVSV-EΔM-tM2e, rVSV-EΔM-HM2e or VSVwt at a multiplicity of infection (MOI) of 0.01. At 12, 24, 48 and 72 hrs after infection, supernatants were collected, and virus titers were measured. The results revealed that VSVwt grew rapidly and reached a titer of 1× 10^9^ TCID_50_ at 24 hours postinfection (Fig. 3D, right panel), while both rVSVΔG/EΔM-tM2e and rVSVΔG/EΔM-HM2e grew slowly and reached a peak titer of 10^8^ TCID_50_ at 48 h postinfection. All rVSVs induced cytopathic effects on Vero E6 cells (Fig. 3D, left panel), indicating the attenuated replication ability of both rVSV-EΔM-tM2e and rVSV-EΔM-HM2e compared to VSVwt.

### 5. rVSV expressing the EΔM-tM2e or EΔM-HM2e fusion protein induced robust anti-M2e and anti-HA IgG antibody responses in immunized mouse serum

We tested whether rVSV-EΔM-tM2e and rVSV-EΔM-HM2e immunization induced immune responses by intramuscularly immunizing Balb/c mice with 1×10^7^ TCID_50_ of rVSV-EΔM-tM2e or rVSV-EΔM-HM2e and administering a booster (5×10^6^ TCID_50_) on Day 21 (Fig. 4A). The sera from immunized mice were collected on Days 20 and 35 postimmunization, and the anti-EboGP antibody responses present in the sera were detected using ELISA. Both the rVSV-EΔM-tM2e- and rVSV-EΔM-HM2e-immunized groups exhibited a high level of anti-EboGP antibodies (Fig. 4B), indicating that immunizations with both rVSV-EΔM-tM2e and rVSV-EΔM-HM2e exhibited similar potency to anti-EboGP immune responses.

**Figure 4:**
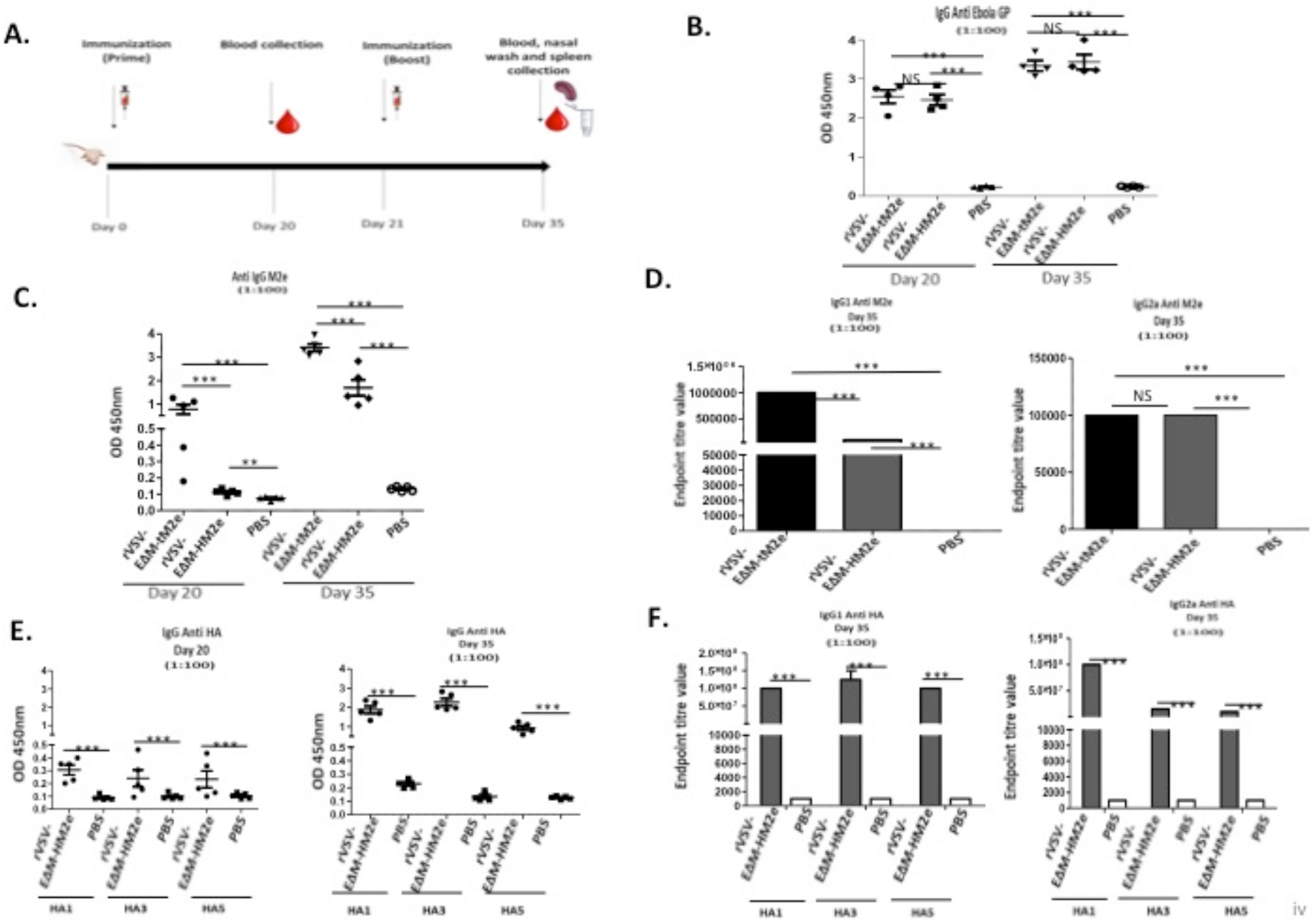
The specific anti-M2e and anti- HA IgG in duced by rVSV-EΔM-tM2e or rVSV-EΔM-HM2e in mice sera. A) The Balb/c mice were injected intramuscularly with 1.0 ×10^7^ TCID_50_ of rVSVEΔM -tM2e or rVSVEΔM-HM2e on Days 0 and 5×10^6^ TCID_50_ on day 21. Serum was collected on Day 20 and 35. ELISA assay was used to detect the levels of anti-Ebola GP IgG (B), Anti-M2e IgG (C), Anti-M2e IgG1, IgG2a (D), anti-HA (HA1, HA3 and HA5) IgG (E) and anti-HA IgG1 and IgG2a (F). Statistical significance between the two groups was determined using an unpaired t-test, and significant p values are represented with asterisks, *≤0.05, **≤0.01. No significance (ns) was shown.

Meanwhile, the anti-M2e antibody level present in the sera from immunized mice was measured. A high titer of anti-M2e IgG antibodies was detected in the sera of the mice immunized with rVSV-EΔM-tM2e, especially after the booster immunization (Fig. 4C). As expected, the anti-M2e IgG level in the rVSV-EΔM-HM2e-immunized mice was 2-3-fold lower than that in the rVSV-EΔM-tM2e-immunized mice (Fig. 4C). We quantitated the anti-M2e IgG subsets present in the total IgG titer by further analyzing the Th1-dependent antibody (IgG2a) and Th2-dependent antibody (IgG1) immune responses. Interestingly, while both rVSV-EΔM-tM2e and rVSV-EΔM-HM2e induced similar levels of anti-M2e IgG2a antibodies (Fig. 4D, left panel), the anti-M2e IgG1 level in the sera of rVSV-EΔM-tM2e-immunized mice was significantly higher than that in the sera of rVSV-EΔM-HM2e-immunized mice (Fig. 4D, right panel).

The anti-HA (including H1, H3 and H5) antibody titers in rVSV-EΔM-HM2e-immunized mice were also evaluated. The results revealed that rVSV-EΔM-HM2e immunization induced robust anti-HA-specific IgG against recombinant HA derived from H1N1, H3N2 and H5N1 (Fig. 4E). Meanwhile, analyses of IgG subsets revealed that rVSV-EΔM-HM2e induced similar levels of anti-HA IgG1 against H1, H3 and H5 but lower levels of Ig2a against H3 and H5 than those against H1 (Fig. 4F). Taken together, rVSV-EΔM-M2e or rVSV-EΔM-HM2e immunization induced robust production of specific anti-M2e and/or anti-HA IgG antibodies in mice.

### 6. Serological and mucosal IgA and cellular immune responses elicited by rVSV expressing EΔM-tM2e or EΔM-HM2e

As shown in previous studies, higher serological and mucosal IgA responses are associated with a better flu disease prognosis and decreased influenza transmission^46, 47, 48, 49^. Therefore, vaccine-induced IgA antibodies are an important immune correlate during influenza vaccination. The levels of anti-M2e or anti-HA IgA in the serum and nasal wash of immunized mice were determined to evaluate the serological and mucosal IgA response to rVSV-EΔM-tM2e or rVSV-EΔM-HM2e. Importantly, high titers of anti-M2e and anti-HA IgA antibodies were detected in the sera of rVSV-EΔM-tM2e- and rVSV-EΔM-HM2e-immunized mice, respectively (Fig. 5A and B). Notably, serological anti-HA titers induced by rVSV-EΔM-HM2e were higher against HA1 than HA3 and HA5. Similarly, rVSV-EΔM-tM2e or rVSV-EΔM-HM2e strongly induced the production of mucosal anti-M2e and anti-HA5 IgA antibodies (Fig. 5C and D). These results indicated the presence of M2e and HA IgA antibodies in the respiratory tracts of the immunized mice.

**Figure 5:**
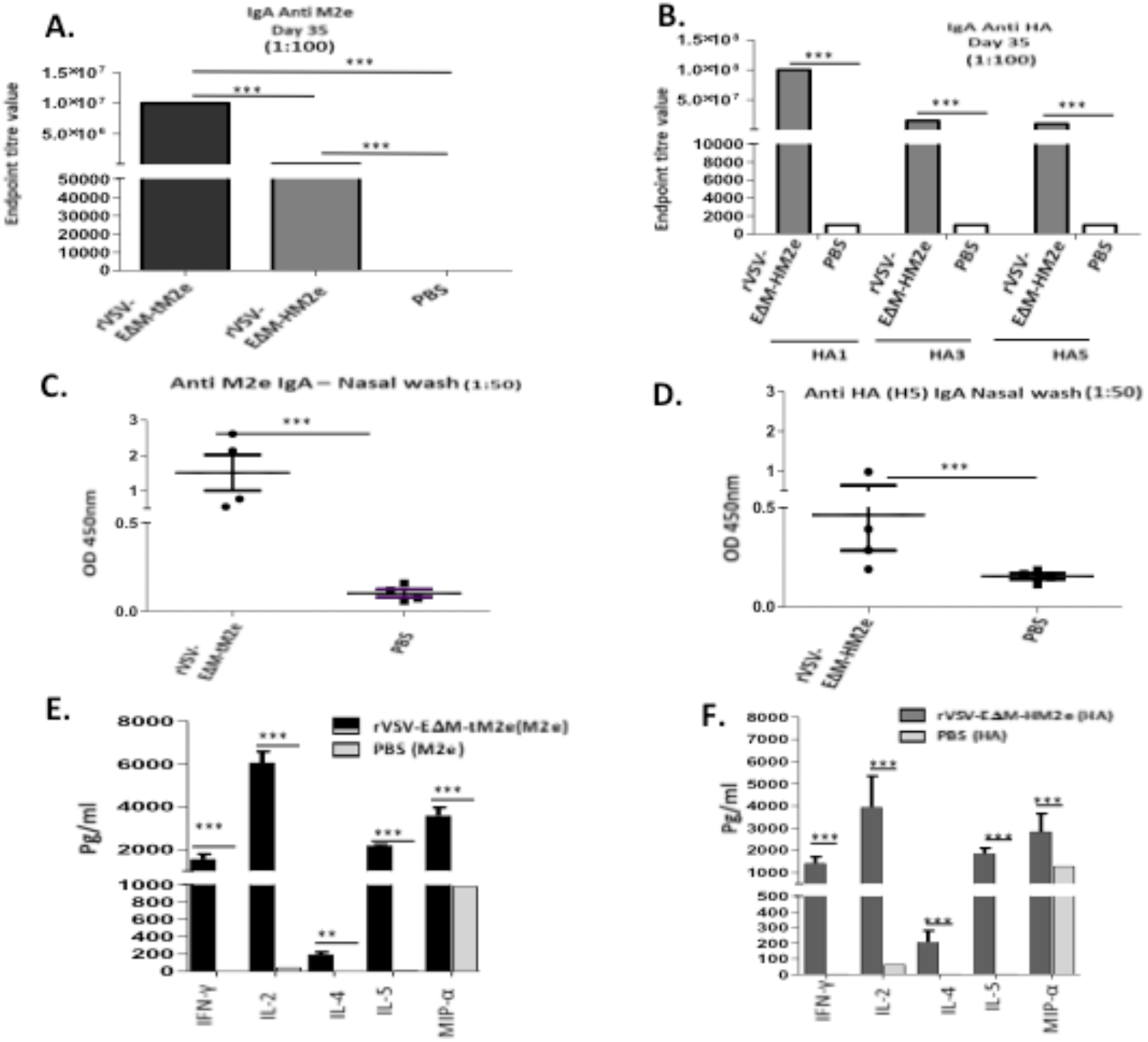
The rVSV EΔM-tM2e or EΔM-HM2e induced specific serological and mucosal Anti-M2e or anti-HA IgA antibodis and mediated cell immune responses. A) the levels of Anti-M2e IgA antibody in mice sera. B) The levels of anti-HA (H1, H3 or H5) IgA antibody in mice sera. C). The levels of Anti-M2e IgA in the nasal wash. D) The levels of IgA anti-HA (5) in the nasal wash. E-F) Splenocytes were isolated from the rVSV/EΔM-tM2e or rVSV/EΔM-HM2e immunized mice and were stimulated with Influenza M2e peptide or HA5 recombinant protein. The release of cytokines and chemokines in the supernatants of cell cultures were quantified with MSD V-plex kit mouse cytokine kit and counted in the MAGPIX instrument. Statistical significance between the two groups was determined using an unpaired t-test, and significant p values are represented with asterisks, *≤0.05, **≤0.01. No significance (ns) was not shown.

Finally, we evaluated the cell-mediated immune responses in immunized mice. For this experiment, splenocytes isolated from rVSV-EΔM-tM2e- or rVSV-EΔM-HM2e-immunized mice were cultured and stimulated with M2e peptides or HA peptides, and the released cytokines and chemokines were quantified using an MSD V-plex mouse cytokine kit, as described in the Materials and Methods. Our results (Fig. 5. E, F) revealed that both rVSV-EΔM-tM2e- and rVSV-EΔM-HM2e-immunized mouse splenocytes expressed significantly higher levels of interferon (IFN)-γ and interleukin (IL)-2, which are involved predominantly in the cellular immune response. Meanwhile, a moderate increase in IL-4 levels and abundant production of IL-5 that may be linked to tissue protection were detected. Immunization using the two vaccine candidates also resulted in the production of macrophage inflammatory protein-1 (MIP-1α).

Overall, rVSV-EΔM-tM2e and rVSV-EΔM-HM2e vaccination resulted in robust vaccine-specific B cell and T cell immune responses.

### 7. Immunization with either rVSV-EΔM-tM2e or rVSV-EΔM-HM2e protects mice from lethal H1N1 and H3N2 influenza virus challenge

We determined whether rVSV-EΔM-tM2e- or rVSV-EΔM-HM2e-induced immune responses protected against lethal influenza virus infection by challenging rVSV-EΔM-tM2e- or rVSV-EΔM-HM2e-immunized mice with mouse-adapted strain A/Puerto Rico/8/34 (H1N1) or H3N2 virus (Table 1). After the intramuscular administration of rVSV-EΔM-tM2e- or rVSV-EΔM-HM2e (primed with 1×10^7^ TCID_50_ and boosted with 5×10^6^ TCID_50_) or PBS, mice were challenged with a low dose (7×10^2^ PFU) of H1N1. The mice vaccinated with PBS showed substantial weight loss until Day 7 before recovering and had a 40% survival rate (Fig. 6A). However, both the rVSV-EΔM-HM2e- and rVSV-EΔM-HM2e-immunized groups showed 100% protection and a slight loss of body weight until Days 5-7 but recovered quickly (Fig. 6A). We then increased the challenge dose of H1N1 to 1.4×10^3^ PFU. The mice vaccinated with PBS experienced morbidity after challenge, as evidenced by a 15 to 30% loss of body mass (Fig. 6B). Interestingly, intramuscular immunization with rVSV-EΔM-tM2e still produced 100% protection in mice, while mice immunized with rVSV-EΔM-HM2e only exhibited 25% protection. We also tested whether the intranasal administration of rVSV-EΔM-tM2e would protect mice from influenza virus challenge. We immunized mice intranasally with a lower dose of rVSV-EΔM-tM2e (1×10^6^ TCID_50_) followed by a booster immunization with 1×10^3^ TCID_50_ of rVSV-EΔM-tM2e (Table 1) and challenge with H1N1 (2.1×10^3^ PFU). The results clearly showed that nasal immunization induced 100% protection, despite the lower dosages of primary and booster immunizations (Fig. 6B).

**Table 1.**

| Experiment Number | Vaccine candidates | Mode of administration | Dosage and day of vaccination (Prime)<br>Day 0<br>(TCID<sub>50</sub>/100µl-Intramuscular)<br>(TCID<sub>50</sub>/50µl-Intranasal) | Dosage and day of vaccination (Boost)<br>Day 14<br>TCID<sub>50</sub>/100µl-Intramuscular)<br>(TCID<sub>50</sub>/50µl-Intranasal) | Viral Challenge<br>Day 28<br>(PFU/50µl- Intranasal) |  |
| --- | --- | --- | --- | --- | --- | --- |
|  |  |  |  |  | H1N1 | H3N2 |
| A | rVSV-EΔM-tM2e (n=5) | Intramuscular | 1x10<sup>7</sup> | 5x10<sup>6</sup> | 700 | NA |
|  | rVSV-EΔM-HM2e (n=5) | Intramuscular | 1x10<sup>7</sup> | 5x10<sup>6</sup> | 700 | NA |
|  | PBS (n=5) | Intramuscular | NA | NA | 700 | NA |
| B | rVSV-EΔM-tM2e (n=4) | Intramuscular | 1x10<sup>7</sup> | 5x10<sup>6</sup> | 2100 | NA |
|  | rVSV-EΔM-tM2e (n=4) | Intranasal | 1x10<sup>6</sup> | 1x10<sup>3</sup> | 2100 | NA |
|  | rVSV-EΔM-HM2e (n=4) | Intramuscular | 1x10<sup>7</sup> | 5x10<sup>6</sup> | 2100 | NA |
|  | PBS (n=4) | Intranasal | NA | NA | 2100 | NA |
| C | rVSV-EΔM-tM2e (n=5) | Intramuscular | 1x10<sup>7</sup> | 5x10<sup>6</sup> | NA | 14000 |
|  | rVSV-EΔM-HM2e (n=5) | Intramuscular | 1x10<sup>7</sup> | 5x10<sup>6</sup> | NA | 14000 |
|  | PBS (n=5) | Intramuscular | NA | NA | NA | 14000 |

**Figure 6:**
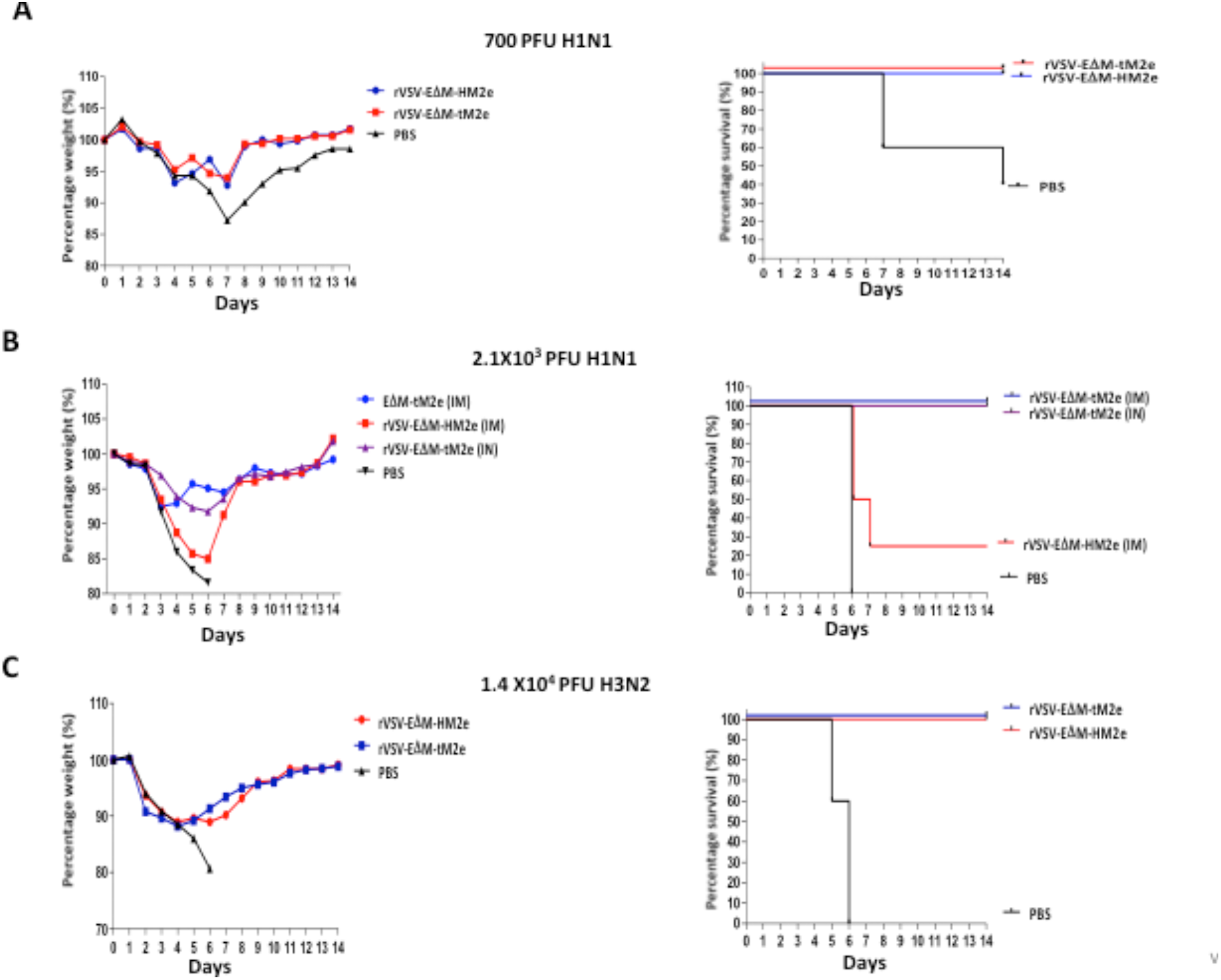
Mice immunized with rVSV-EΔM-tM2e or rVSV-EΔM-HM2e were protected against challenge with H1N1 or H3N2 influenza virus strains. The Balb/c mice were injected intramuscularly (im) with 1×10^7^ TCID_50_ of rVSV/EΔM -tM2e, rVSV/EΔM-HM2e or PBS Days 0 and 5×10^6^ TCID_50_ on Day 14. Alternatively, mice were intranasally (in) vaccinated with 1×10^6^ TCID_50_ of rVSV/EΔM -tM2e on Day 0 and 1×10^3^ TCID_50_ of rVSV/EΔM-tM2e on Day 14. On day 28 post-immunization, all the mice were challenged with 700 PFU (A) or 2.1×10^3^ PFU (B) of H1N1, or 1.4×104 PFU (C) of H3N2. Weight loss or gain of the mice were monitored daily for 2 weeks (A,B,C Left panel) and the survive rates of mice infected with H1N1 or H3N2 was shown at the Right panel.

The mice immunized intramuscularly with rVSV-EΔM-tM2e or rVSV-EΔM-HM2e (primed with a dose of 1×10^7^ TCID_50_ and boosted with 5×10^6^ TCID_50_) were infected with 1.4×10^4^ PFU of mouse-adapted H3N2 virus to investigate the protection provided by our VSV-based vaccine candidates against another subtype (H3N2) of influenza virus (Table 1). Mice vaccinated with rVSV-EΔM-tM2e or rVSV-EΔM-HM2e showed slight weight loss until Day 5 and were fully protected (100% survival rate), while the infected control (PBS) group did not survive beyond Day 6 (Fig. 6C). All these results provided evidence that immunization with rVSV-EΔM-tM2e and rVSV-EΔM-HM2e efficiently protected mice from lethal H1N1 and H3N2 influenza challenges, while rVSV-EΔM-tM2e produced more effective protection against the higher dose of H1N1 challenge.

## Discussion

Despite the progress achieved in developing a universal vaccine for influenza viral infection, an FDA-approved universal vaccine for influenza viral infection is still unavailable. In this study, we generated PVP and rVSV-based vaccine candidates expressing the ectodomain of influenza matrix protein (M2e) and/or hemagglutinin stalk regions (HAcs) that were fused with the DC-targeting domain of EboGP (EΔM) and revealed their abilities to efficiently elicit anti-influenza immune responses and protect against different strains of influenza virus.

Recent advances in influenza virus vaccine research have shown that targeting the highly conserved epitopes in the HA stalk domain or M2e of influenza virus is a promising approach for developing a universal vaccine ^13 50, 51, 52^. However, methods to enhance the antigenicity of these large polypeptides are still an important issue to be addressed. According to our recent study, the replacement of the mucin-like domain (MLD) of EboGP with heterologous large polypeptides still maintains the ability of EboGP to target human macrophages/dendritic cells (DCs) and induce robust immune responses against the inserted polypeptides ^53^. Therefore, in this study, we fused four copies of M2e from human (2 copies), avian (one copy) and swine (one copy) influenza strains with EΔM or M2e (from human influenza virus) and influenza HAcs derived from H5N1 (aa 156) ^54^ with EΔM to generate EΔM-tM2e or EΔM-HM2e fusion proteins, respectively (Fig. 1A-C), with increased immunogenicity. Because the mucin-like domain is located at the apex of the sides of the EboGP monomer ^34, 55^, the inserted heterologous polypeptides, such as HM2e and tM2e, are also expected to be exposed to the apex of the sides of the EΔM-HM2e and EΔM-M2e fusion proteins and may not interfere with the cell targeting and entry of EΔM. Indeed, our study showed that EΔM-HM2e- and EΔM-tM2e-PVPs exhibited a strong ability to target and enter DCs and macrophages compared to native HA/NA/M2-PVPs (Fig. 1F-G). This finding also correlated with our recently published data revealing that the incorporation of the EΔM-V3 fusion protein or EboGP into HIV Env pseudotyped VLPs facilitated DC targeting compared to HIV-Env PVPs, indicating that EΔM accommodates large polypeptides without losing its DC and macrophage targeting ability ^28, 53^.

As the presence of EboGP or EΔM-V3 fusing protein in either PVP- or rVSV-based vaccine candidates substantially enhances their immunogenicity ^28, 34^, we also tested whether EΔM-M2e or EΔM-HM2e-PVPs displayed strong immunogenicity. For this experiment, the mice were immunized intramuscularly with EΔM-M2e- or EΔM-HM2e-PVPs, and the results clearly showed that the anti-M2e and/or anti-HA antibody titers in EΔM-tM2e- or EΔM-HM2e-PVP-immunized mice were significantly higher than those in HA/NA/M2-PVP-immunized mice (Fig. 2C-E). The anti-p24 antibody titer in EΔM-tM2e-PVP- or EΔM-HM2e-PVP-immunized mice was also higher than that in HA/NA/M2-PVP-immunized mice (Fig. 2F). Moreover, our results clearly showed that EΔM-M2e-PVPs resulted in remarkably higher anti-M2 antibody levels that exceeded the anti-M2 antibody titer produced following EΔM-HM2e-PVP immunization (Fig. 2C and D). Overall, the aforementioned observations provide evidence to support our hypothesis that the presence of EΔM-tM2e or EΔM-HM2e on the surface of PVPs induces efficient antibody responses to M2 and HA.

The rVSV vector system as a vaccine platform has attracted global attention as a vaccine delivery system for multiple viral proteins that induces strong humoral and cell-mediated immune responses against viral proteins ^53, 56, 57, 58 59^. Interestingly, a recent study showed that rVSV expressing HA stalk as an influenza vaccine confers rapid protection against different H5 influenza strains ^18^. In this study, we generated rVSVs encoding EΔM-tM2e or EΔM-HM2e in place of VSV-G and found that the replication of rVSV-EΔM-tM2e or rVSV-EΔM-HM2e was significantly attenuated compared with VSVwt (Fig. D). This observation is important for the safety of rVSV-based vaccines. In the animal study, a single dose of rVSV immunization induced relatively high levels of anti-M2e and anti-HA antibody responses on Day 20. Following the booster immunization, both rVSV-EΔM-tM2e and rVSV-EΔM-HM2e induced robust anti-influenza HA and M2e IgG production, including IgG2a and IgG1, and significantly higher levels of anti-influenza IgA antibodies in both the sera and nasal mucosa of mice (Figs. 4 and 5). Furthermore, the splenocytes of rVSV-EΔM-tM2e- and rVSV-EΔM-HM2e-immunized mice released large amounts of MIP-1α, IL-2, IL-5, IL-6, IL-10 and IFN-γ upon stimulation with M2e peptide or HA peptide, indicating that immunization with these rVSV vaccine candidates also induced significantly higher levels of cell-mediated immunity (Fig. 5E, F). Unfortunately, we still do not know whether these vaccine candidates might induce stronger cell-mediated immunity than the wild-type influenza HA or M2 protein due to a lack of rVSV expressing the wild-type influenza HA or M2 protein in our experimental system.

Another interesting observation was that rVSV-EΔM-tM2e elucidated a remarkably higher level of the anti-M2e antibody than rVSV-EΔM-HM2e (Fig. 4). This result was consistent with the data from EΔM-tM2e-PVPs (Fig. 2C). M2e has low immunogenicity due to its small size (aa 24) and small number of copies in the virus. We increased its immunogenicity by creating PVPs or rVSV containing four M2e tandem repeats that increase the M2e copy number on the surface of viral particles and/or the cells. This strategy resulted in significant increases in the production of M2e-specific antibodies and cell-mediated immune response. The intranasal and intramuscular immunizations of rVSV-EΔM-tM2e provided 100% protection against both H1N1 and H3N2 challenges (Fig. 6). Thus, using EboGP DC-targeting domain (EΔM) fusion protein technology ^34^, we can insert multiple copies of large polypeptides in EΔM to induce significantly stronger immune responses against targeted polypeptides. Additionally, this study provides evidence for the efficacy of the rVSV-EΔM-tM2e vaccine candidate and for M2e-based vaccines as a universal vaccine approach.

Our study also showed that rVSV-EΔM-HM2e induced high levels of antibody responses to HA from different influenza strains, including H1N1, H3N2 and H5N1 (Fig. 4E and Fig. 5B), indicating its broad anti-HA responses. The rVSV-HAcs-M2e vaccine immunization also provided 100% protection against challenge with a lethal dose of H3N1 and partial (25%) protection against lethal H1N1 challenge (Fig. 6). Based on these results, the rVSV-tM2e vaccine candidate appears to provide broader or more efficient protection than rVSV-HM2e. A universal influenza vaccine should be effective against all influenza viruses, regardless of any antigenic mutation or HA/NA subtypes. Further studies are required to investigate the protective effects of these candidate vaccines on other strains of influenza virus and on nonhuman primates. However, the results presented in this study have provided strong evidence for a universal influenza vaccine platform.

Overall, this study is the first to show that infusion of the influenza HA stalk region and/or conserved M2e with EΔM elicited robust influenza immune responses and protected against different strains of influenza virus. The novelty of this study is the use of EboGP-based DC-targeting domain (EΔM) fusion protein technology to present the conserved antigens of influenza virus (HA stalk regions and conserved M2e region) and induce a stronger immune response against influenza virus infections. This study also provides convincing evidence for the EboGP-based DC-targeting domain (EΔM) fusion technology that can be broadly used to develop vaccine strategies that protect against other emergent and re-emergent infectious diseases.

## Materials and methods

### Plasmid constructions

In this study, the mucin-like domain (MLD) deletion-containing Ebola glycoprotein (EboGP) plasmid (pCAGGS-EboGPΔM) were described previously ^28^. To construct pCAGGS-EΔM-tM2e and pCAGGS-EΔM-HM2e plasmids, we designed and synthesized genes encoding for HAcsM2e (aa 179) or tM2e (aa 92) (Fig. 1A) and inserted these genes into pCAGGS-EΔM through *ApaI* and *XbaI* sites (Figure 1B). To construct rVSV vector expressing EΔM-tM2e or EΔM-HM2e plasmids, we inserted the PCR amplified cDNA encoding a full-EΔM-tM2e or EΔM-HM2e at the *MluI* and *SphI* sites in a VSV-G deleted rVSV vector, as described previously ^34^ (Figure 3A).

The pCAGGS-HA, -NA, and -M2 plasmids and the HIV-1 RT/IN/Env tri-defective proviral plasmid containing Gaussia gene (ΔRI/ΔE/Gluc) were described previously^37 28^

### Cells, antibodies, and chemicals

Human embryonic kidney 293T, Vero E6 cell lines were cultured in DMEM supplemented with 10% fetal bovine serum (FBS). To obtain MDMs or MDDCs, monocytes were isolated from human peripheral blood mononuclear cells (hPBMCs) by sedimentation on a Ficoll (Lymphoprep; Axis-Shield) gradient and treated with macrophage colony stimulator (M-CSF) or granulocyte-macrophage-stimulating factor (GM-CSF) and IL-4 (R&D system) respectively for 7 days. The M2e monoclonal antibody and HA polyclonal antibody were obtained from Santa Cruz Biotechnology (14C2: sc-32238) and Alpha Diagnostic (HA2H012-A) respectively. Ebola GP monoclonal antibody (mAb) 42/3.7 was kindly given by Dr. A Takada, Hokkaido University, Japan ^60^. M2e peptide was purchased from Genescript (RP20206) and HA peptide was synthesized by Shangai Royobiotech (19CL00157). The recombinant HA from H1N1 (11684-V08H), H3N2 (40494-V08B) and H5N1 (40160-V08B1) were obtained from Sino Biological.

### Production and characterization of PVPs or rVSV containing EΔM-tM2e and EΔM-HM2e

To produce EΔM-tM2e or EΔM-HM2e pseudotyped viral particles (PVPs) for *in vitro* study of viral entry, 293T cells were co-transfected pCAGGS-EΔM-tM2e or pCAGGS-EΔM-HM2e with Δ8.2 and ΔRI/ΔE/Gluc+. Meanwhile, a native HA/NA/M2 PVPs was produced by co-transfected pCAGGS-HA, -NA, and -M2 plasmids with Δ8.2 and ΔRI/ΔE/Gluc+ and used as control. For *in vivo* immunization experiment, 293T cells were transfected with the above plasmids except for ΔRI/ΔE/Gluc+. After 48 h of post-transfection, the PVPs particles were pelleted by ultracentrifugation at 35000rpm. The virus stocks were quantified by using HIV p24 ELISA assay and kept at -80°C.

By using reverse genetics technology ^44^, the rVSV-EΔM-tM2e or rVSV-EΔM-HM2e vector was transfected into a mix of VeroE6/293T cells together with VSV accessory plasmids encoding for P, L, N, and T7 promoter plasmid. Following the primary transfection, the supernatant containing the recovered rVSV-EΔM-tM2e or rVSV-EΔM-HM2e were amplified in Vero E6 cells. Produced VSV was concentrated by ultracentrifugation, titrated in VeroE6 and used for mice immunization experiments.

To detect the expression and incorporation of M2e, HAcs and other viral proteins in cells, the transfected cells and PVPs or rVSV were lysed and analyzed by SDS-PAGE and WB with Anti-M2e (14C2), anti-HA, EBOV GP MAb 42/3.7 or anti HIVp24 antibodies, respectively.

### The *Gaussia* luciferase (GLuc) assay

To test the entry ability of PVPs to MDMs and MDDCs, an equal amount (adjusted by P24) of EΔM-, EΔM-HM2e-, EΔM-tM2e-GLuc+ PVPs were used to infect MDMs and MDDCs. The *Gaussia* luciferase (GLuc) assay was done at various times of infection by collecting supernatants from the cell cultures. A 10µl of sample was mixed with a 50µl portion of GAR-1 reagent (targeting Systems) and then measured in the luminometer (Promega, USA)^36^.

### Immunofluorescence assay

As previously described ^45^, Vero E6 cells were grown on a glass coverslip (12 mm^2^) in a 24-well plate and infected with rVSV/EΔM-HM2e or rVSV/EΔM-tM2e for 48 hours. After infection, cells on the coverslip were fixed with PBS in 4% paraformaldehyde for 5 minutes, permeabilized with 0.2% Triton X-100 in PBS. The cells were then incubated in primary antibodies specific for M2e or HA followed by corresponding secondary FITC-conjugated secondary antibodies. The cells were viewed under a computerized Axiovert 200 fluorescence microscopy (Becto Deckson).

### Mice immunization experiment

Female BALB/c mice aged 4–6 weeks used in this study were obtained from the Central Animal Care Facility, the University of Manitoba (with the animal study protocol approval No. 12-017). For PVPs immunization, the mice were divided into four per group. each group was immunized subcutaneously by injection with EΔM-tM2e, EΔM-HM2e, native HA/NA/M2 100 ng (adjusted by HIV p24) of PVPs in 100 µl endotoxin-free PBS or PBS only on days 0, 28 and 56. Blood samples were collected on Day 63 and the mice were sacrificed. For rVSV immunization, mice (five for each group) were immunized intramuscularly with 1×10^7^ TCID_50_ rVSV-EΔM-tM2e or rVSV-EΔM-HM2e and boosted on day 21 with 5×10^6^ TCID_50_ of VSV and sacrificed on day 35. Blood samples, nasal wash and splenocytes were collected on day 35.

For influenza viral challenge in mice, the mouse-adapted strain of A/Puerto Rico/8/34 (H1N1) or H3N2 virus strains were used. Each group of mice (5 for each group) were intramuscularly immunized with 1×10^7^ TCID_50_ of rVSV-EΔM-tM2e, rVSV-EΔM-HM2e or PBS and boosted with 5×10^6^ TCID_50_ of VSV or PBS on day 21. Meanwhile, One group of mice were intranasal immunization with 1×10^6^ TCID_50_ of EΔM-tM2e and was boosted with 1×10^3^ TCID_50_ on day 21. After two weeks of the final immunization, the challenge was done by infecting the mice intranasally with 2.1×10^3^ PFU or 7×10^2^ PFU H1N1, or 1.4×10^4^ PFU of H3N2, while mice injected with PBS were challenged as a negative control. Weight loss or gain of the mice were monitored daily for 2 weeks after the challenge.

### Mouse serum and nasal wash antibodies measurements by Enzyme-linked Immunosorbent Assay (ELISA)

To determine influenza HA and M2 specific antibodies in mice sera, ELISA plates (NUNC Maxisorp, Thermo Scientific) were coated with 100 µl of M2e peptide or rHA (H1, H3 or H5) proteins (1µg or 0.5 µg/ml respectively) in a coupling buffer (0.05M carbonate-bicarbonate of pH 9.6) overnight at 4°C. To measure the HIV Gag-specific or EboGP-specific antibody in sera, the plate was coated with HIV-1 IIIB p24 recombinant protein (0.5µg/ml) or recombinant glycoprotein (0.5µg/ml) as described previously^53^. The plates were washed twice after the incubation with 1X PBST and blocked with blocking buffer (1% BSA in PBS) at 37°C for 1 hr. The serum samples were diluted with primary antibody diluent (1:100-1:10^9^), then 100µl of the diluted mouse serum samples were added into each well of the plates and incubated for 2 hrs at 37°C. Dishes were immediately washed five times, and 100 µl peroxidase-conjugated goat anti-mouse immunoglobulin G (IgG) (GE Healthcare), IgG1, IgG2a or IgA secondary antibodies were added and incubated for 1 h at 37°C. The plates were washed five times, and 3’,3’,5’,5’ Tetramethylbenzidine (TMB) (Mandel Scientific) was added and incubated for 15 min at room temperature in a dark room. To stop the reaction, 100 µl of 1N HCl was added to each well, and the absorbance was measured at 450nm optical density ^18^.

### Cytokine detection

Splenocytes from immunized mice have collected aseptically and placed into the cell strainer and were mashed through the cell strainer inside a sterile 50 ml tube using the plunger end of the syringe to make single-cell suspensions. Red blood cells were removed using Ammonium-Chloride-Potassium (ACK) buffer, and the suspended cell was cultured in 48-well plates at a density of 2×10^6^/125µl with DMEM containing either M2 or HA peptide (1 or 2 µg/peptide/ml) respectively. After 3 days of culturing, supernatants were collected and stored at - 70°C for a cytokine detection assay. Cytokine (IFN-g, IL-2, IL-4, IL-5, and MIP-1a) levels were measured in supernatants using the 8-plex mouse MSD V-plex kit (Mesoscale Discovery, USA) following the manufacturer procedure.

### Statistics

Statistical analysis of cytokine levels, including the results of GLuc assay, influenza M2, HA and ELISA, and various cytokine/chemokines, were performed using the unpaired t-test (considered significant at P≥0.05) by GraphPad Prism 5.01 software.

## Acknowledgements

We thank Mahmud-ur Rashid for technical assistance. Titus Olukitibi is a recipient of the University Manitoba Graduate scholarship. This work was supported by the Manitoba Research Chair Award from the Research Manitoba (RM) and Canadian HIV Vaccine Initiative (CHVI) Bridge Funding (201309OCB) to X-J.Y.

## References

1. Nickol ME, Kindrachuk J. A year of terror and a century of reflection: perspectives on the great influenza pandemic of 1918-1919. BMC infectious diseases 2019, 19(1): 117.

2. Kedzierska K, Van de Sandt C, Short KR. Back to the future: lessons learned from the 1918 influenza pandemic. Frontiers in Cellular and Infection Microbiology 2018, 8: 343.

3. Valkenburg SA, Mallajosyula VVA, Li OTW, Chin AWH, Carnell G, Temperton N, et al. Stalking influenza by vaccination with pre-fusion headless HA mini-stem. Scientific reports 2016, 6(22666): 1–11.

4. Francis ME, King ML, Kelvin AA. Back to the Future for Influenza Preimmunity “Looking Back at Influenza Virus History to Infer the Outcome of Future Infections. Viruses 2019, 11(2): 122.

5. Tong S, Zhu X, Li Y, Shi M, Zhang J, Bourgeois M, et al. New world bats harbor diverse influenza A viruses. PLoS pathogens 2013, 9(10): e1003657.

6. Scorza FB, Tsvetnitsky V, Donnelly JJ. Universal influenza vaccines: Shifting to better vaccines. Vaccine 2016, 34(26): 2926–2933.

7. Tsilibary EP, Charonis SA, Georgopoulos AP. Vaccines for Influenza. Vaccines 2021, 9, 47. 2021.

8. Krammer F, Palese P. Influenza virus hemagglutinin stalk-based antibodies and vaccines. Current opinion in virology 2013, 3(5): 521–530.

9. Mullarkey CE, Bailey MJ, Golubeva DA, Tan GS, Nachbagauer R, He W, et al. Broadly neutralizing hemagglutinin stalk-specific antibodies induce potent phagocytosis of immune complexes by neutrophils in an Fc-dependent manner. MBio 2016, 7(5): e01624–01616.

10. Cummings JF, Guerrero ML, Moon JE, Waterman P, Nielsen RK, Jefferson S, et al. Safety and immunogenicity of a plant-produced recombinant monomer hemagglutinin-based influenza vaccine derived from influenza A (H1N1) pdm09 virus: a Phase 1 dose-escalation study in healthy adults. Vaccine 2014, 32(19): 2251–2259.

11. Uranowska K, Tyborowska J, Jurek A, Szewczyk Ba, Gromadzka B. Hemagglutinin stalk domain from H5N1 strain as a potentially universal antigen. Acta Biochimica Polonica 2014, 61(3).

12. Von Holle TA, Moody MA. Influenza and antibody-dependent cellular cytotoxicity. Frontiers in immunology 2019, 10: 1457.

13. Mezhenskaya D, Isakova-Sivak I, Rudenko L. M2e-based universal influenza vaccines: a historical overview and new approaches to development. Journal of biomedical science 2019, 26(1): 1–15.

14. Tsybalova LM, Stepanova LA, Shuklina MA, Mardanova ES, Kotlyarov RY, Potapchuk MV, et al. Combination of M2e peptide with stalk HA epitopes of influenza A virus enhances protective properties of recombinant vaccine. PloS one 2018, 13(8): e0201429.

15. Krammer F, Palese P, Steel J. Advances in universal influenza virus vaccine design and antibody mediated therapies based on conserved regions of the hemagglutinin. Influenza Pathogenesis and Control-Volume II. Springer, 2014, pp 301–321.

16. Turley CB, Rupp RE, Johnson C, Taylor DN, Wolfson J, Tussey L, et al. Safety and immunogenicity of a recombinant M2e-”flagellin influenza vaccine (STF2. 4xM2e) in healthy adults. Vaccine 2011, 29(32): 5145–5152.

17. Li X, Pushko P, Tretyakova I. Recombinant hemagglutinin and virus-like particle vaccines for H7N9 influenza virus. Journal of vaccines & vaccination 2015, 6(3).

18. Furuyama W, Reynolds P, Haddock E, Meade-White K, Le MQ, Kawaoka Y, et al. A single dose of a vesicular stomatitis virus-based influenza vaccine confers rapid protection against H5 viruses from different clades. NPJ vaccines 2020, 5(1): 1–10.

19. Nachbagauer R, Feser J, Naficy A, Bernstein DI, Guptill J, Walter EB, et al. A chimeric hemagglutinin-based universal influenza virus vaccine approach induces broad and long-lasting immunity in a randomized, placebo-controlled phase I trial. Nature medicine 2020, 27(1): 106–114.

20. Brandao JG, Scheper RJ, Lougheed SaM, Curiel DT, Tillman BW, Gerritsen WR, et al. CD40-targeted adenoviral gene transfer to dendritic cells through the use of a novel bispecific single-chain Fv antibody enhances cytotoxic T cell activation. Vaccine 2003, 21(19-20): 2268–2272.

21. Querec T, Bennouna S, Alkan S, Laouar Y, Gorden K, Flavell R, et al. Yellow fever vaccine YF-17D activates multiple dendritic cell subsets via TLR2, 7, 8, and 9 to stimulate polyvalent immunity. Journal of Experimental Medicine 2006, 203(2): 413–424.

22. Banchereau J, Palucka AK. Dendritic cells as therapeutic vaccines against cancer. Nature Reviews Immunology 2005, 5(4): 296.

23. Gudjonsson A, LysÃ©n A, Balan S, Sundvold-Gjerstad V, Arnold-Schrauf C, Richter L, et al. Targeting influenza virus hemagglutinin to Xcr1+ dendritic cells in the absence of receptor-mediated endocytosis enhances protective antibody responses. The Journal of Immunology 2017, 198(7): 2785–2795.

24. Park H-Y, Tan PS, Kavishna R, Ker A, Lu J, Chan CEZ, et al. Enhancing vaccine antibody responses by targeting Clec9A on dendritic cells. NPJ vaccines 2017, 2(1): 31.

25. Yamasaki S, Shimizu K, Kometani K, Sakurai M, Kawamura M, Fujii SI. In vivo dendritic cell targeting cellular vaccine induces CD4(+) Tfh cell-dependent antibody against influenza virus. 2016(2045–2322 (Electronic)).

26. Martinez O, Leung LW, Basler CF. The role of antigen-presenting cells in filoviral hemorrhagic fever: gaps in current knowledge. Antiviral research 2012, 93(3): 416–428.

27. Martinez O, Johnson JC, Honko A, Yen B, Shabman RS, Hensley LE, et al. Ebola virus exploits a monocyte differentiation program to promote its entry. Journal of virology 2013, 87(7): 3801–3814.

28. Ao Z, Wang L, Mendoza EJ, Cheng K, Zhu W, Cohen EA, et al. Incorporation of Ebola glycoprotein into HIV particles facilitates dendritic cell and macrophage targeting and enhances HIV-specific immune responses. PloS one 2019, 14(5): e0216949.

29. Olukitibi TA, Ao Z, Mahmoudi M, Kobinger GA, Yao X. Dendritic Cells/Macrophages-Targeting Feature of Ebola Glycoprotein and its Potential as Immunological Facilitator for Antiviral Vaccine Approach. Microorganisms 2019, 7(10): 402–425.

30. Tran EE, Simmons JA, Bartesaghi A, Shoemaker CJ, Nelson E, White JM, et al. Spatial localization of the Ebola virus glycoprotein mucin-like domain determined by cryo-electron tomography. Journal of virology 2014, 88(18): 10958–10962.

31. Medina MF, Kobinger GP, Rux J, Gasmi M, Looney DJ, Bates P, et al. Lentiviral vectors pseudotyped with minimal filovirus envelopes increased gene transfer in murine lung. Molecular therapy: the journal of the American Society of Gene Therapy 2003, 8(5): 777–789.

32. Lee JE, Fusco ML, Hessell AJ, Oswald WB, Burton DR, Saphire EO. Structure of the Ebola virus glycoprotein bound to a human survivor antibody. Nature 2008.

33. Lee JE, Fusco ML, Hessell AJ, Oswald WB, Burton DR, Saphire EO. Structure of the Ebola virus glycoprotein bound to a human survivor antibody. Nature 2009.

34. Ao Z-j, Olukitibi T, Wang L-j, Azizi H, Mahmoudi M, Kobinger G, et al. Development and Evaluation of an Ebola Virus Glycoprotein Mucin-Like Domain Replacement System as a New Dendritic Cell-Targeting Vaccine Approach against HIV-1. Journal of virology 2021, 95: Jul 12;95(15):(12): e0236820. PMID: 34011553.

35. Krammer F. The human antibody response to influenza A virus infection and vaccination. Nature Reviews Immunology 2019, 19(6): 383–397.

36. Ao Z, Huang J, Tan X, Wang X, Tian T, Zhang X, et al. Characterization of the single cycle replication of HIV-1 expressing Gaussia Luciferase in human PBMCs, macrophages, and in CD4+ T cell-grafted nude mouse. Journal of virological methods 2016, 228: 95–102.

37. Ao Z, Patel A, Tran K, He X, Fowke K, Coombs K, et al. Characterization of a trypsin-dependent avian influenza H5N1-pseudotyped HIV vector system for high throughput screening of inhibitory molecules. Antiviral research 2008, 79(1): 12–18.

38. Alvarez CP, Lasala Ft, Carrillo J, MuÃ±iz O, CorbÃ-AL Delgado R. C-type lectins DC-SIGN and L-SIGN mediate cellular entry by Ebola virus in cis and in trans. Journal of virology 2002, 76(13): 6841–6844.

39. Ye L, Lin J, Sun Y, Bennouna S, Lo M, Wu Q, et al. Ebola virus-like particles produced in insect cells exhibit dendritic cell stimulating activity and induce neutralizing antibodies. Virology 2006, 351(2): 260–270.

40. Thompson CM, Petiot E, Lennaertz A, Henry O, Kamen AA. Analytical technologies for influenza virus-like particle candidate vaccines: challenges and emerging approaches. Virology journal 2013, 10(1): 1–14.

41. Geisbert TW, Feldmann H. Recombinant vesicular stomatitis virus-”based vaccines against Ebola and Marburg virus infections. The Journal of infectious diseases 2008, 204(uppl_3): S1075–S1081.

42. Geisbert TW, Feldmann H. Recombinant vesicular stomatitis virus-”based vaccines against Ebola and Marburg virus infections. The Journal of infectious diseases 2011, 204(suppl_3): S1075–S1081.

43. Fathi A, Dahlke C, Addo MM. Recombinant vesicular stomatitis virus vector vaccines for WHO blueprint priority pathogens. Human vaccines & immunotherapeutics 2019, 15(10): 2269–2285.

44. Witko SE, Kotash CS, Nowak RM, Johnson JE, Boutilier LA, Melville KJ, et al. An efficient helper-virus-free method for rescue of recombinant paramyxoviruses and rhadoviruses from a cell line suitable for vaccine development. Journal of virological methods 2006, 135(1): 91–101.

45. Ao Z, Fowke KR, Cohen ÃrA, Yao X. Contribution of the C-terminal tri-lysine regions of human immunodeficiency virus type 1 integrase for efficient reverse transcription and viral DNA nuclear import. Retrovirology 2005, 2(1): 1–15.

46. Clements ML, Betts RF, Tierney EL, Murphy BR. Serum and nasal wash antibodies associated with resistance to experimental challenge with influenza A wild-type virus. Journal of clinical microbiology 1986, 24(1): 157–160.

47. Belshe RB, Gruber WC, Mendelman PM, Mehta HB, Mahmood K, Reisinger K, et al. Correlates of immune protection induced by live, attenuated, cold-adapted, trivalent, intranasal influenza virus vaccine. The Journal of infectious diseases 2000, 181(3): 1133–1137.

48. Ambrose CS, Wu X, Jones T, Mallory RM. The role of nasal IgA in children vaccinated with live attenuated influenza vaccine. Vaccine 2012, 30(48): 6794–6801.

49. Abreu RB, Clutter EF, Attari S, Sautto GA, Ross TM. IgA responses following recurrent influenza virus vaccination. Frontiers in immunology 2020, 11: 902.

50. Marzi A, Yoshida R, Miyamoto H, Ishijima M, Suzuki Y, Higuchi M, et al. Protective efficacy of neutralizing monoclonal antibodies in a nonhuman primate model of Ebola hemorrhagic fever. PloS one 2019, 7(4): e36192.

51. Gunn BM, Yu W-H, Karim MM, Brannan JM, Herbert AS, Wec AZ, et al. A role for Fc function in therapeutic monoclonal antibody-mediated protection against Ebola virus. Cell host & microbe 2018, 24(2): 221–233. e225.

52. Konduru K, Shurtleff AC, Bradfute SB, Nakamura S, Bavari S, Kaplan G. Ebolavirus glycoprotein Fc fusion protein protects guinea pigs against lethal challenge. PloS one 2016, 11(9): e0162446.

53. Ao Z, Wang L, Azizi H, Abiola TO, Kobinger G, Yao X. Development and Evaluation of an Ebola Virus Glycoprotein Mucin-Like Domain Replacement System as a New DC-Targeting Vaccine Approach Against HIV-1. Journal of virology 2021: JVI. 02368-02320.

54. Maines TR, Lu XH, Erb SM, Edwards L, Guarner J, Greer PW, et al. Avian influenza (H5N1) viruses isolated from humans in Asia in 2004 exhibit increased virulence in mammals. Journal of virology 2005, 79(18): 11788–11800.

55. Tran EEH, Simmons JA, Bartesaghi A, Shoemaker CJ, Nelson E, White JM, et al. Spatial localization of the Ebola virus glycoprotein mucin-like domain determined by cryo-electron tomography. Journal of virology 2014, 88(18): 10958–10962.

56. Lai C-Y, Strange DP, Wong TAS, Lehrer AT, Verma S. Ebola Virus Glycoprotein Induces an Innate Immune Response In vivo via TLR4. Frontiers in Microbiology, 8(1571).

57. Geisbert TW, Geisbert JB, Leung A, Daddario-DiCaprio KM, Hensley LE, Grolla A, et al. Single-injection vaccine protects nonhuman primates against infection with Marburg virus and three species of Ebola virus. Journal of virology 2009, 83(14): 7296–7304.

58. Sun W, Zheng A, Miller R, Krammer F, Palese P. An inactivated influenza virus vaccine approach to targeting the conserved hemagglutinin stalk and M2e domains. Vaccines 2019, 7(3): 117.

59. Grohskopf LA, Munoz FM. 2018-2019 Recommendations for influenza prevention and treatment in children: an update for pediatric providers. 2018.

60. Takada A, Ebihara H, Feldmann H, Geisbert TW, Kawaoka Y. Epitopes required for antibody-dependent enhancement of Ebola virus infection. The Journal of infectious diseases 2007, 196(Supplement_2): S347–S356.

61. Gauger PC, Vincent AL. Serum virus neutralization assay for detection and quantitation of serum-neutralizing antibodies to influenza A virus in swine. Animal influenza virus. Springer, 2014, pp 313–324.

